# Coordination of brain wide activity dynamics by dopaminergic neurons

**DOI:** 10.1101/042168

**Authors:** Heather K. Decot, Vijay M.K. Namboodiri, Wei Gao, Jenna A. McHenry, Joshua H. Jennings, Sung-Ho Lee, Pranish A. Kantak, Yu-Chieh Jill Kao, Manasmita Das, Ilana B. Witten, Karl Deisseroth, Yen-Yu Ian Shih, Garret D. Stuber

## Abstract

Several neuropsychiatric conditions, such as addiction, schizophrenia, and depression may arise in part from dysregulated activity of ventral tegmental area dopaminergic (TH^VTA^) neurons, as well as from more global maladaptation in neurocircuit function. However, whether TH^VTA^ activity affects large-scale brain-wide function remains unknown. Here, we selectively activated TH^VTA^ neurons in transgenic rats and measured resulting changes in whole-brain activity using stimulus-evoked functional magnetic resonance imaging (fMRI). Selective optogenetic stimulation of TH^VTA^ neurons not only enhanced cerebral blood volume (CBV) signals in striatal target regions in a dopamine receptor dependent fashion, but also engaged many additional anatomically defined regions throughout the brain. In addition, repeated pairing of TH^VTA^ neuronal activity with forepaw stimulation, produced an expanded brain-wide sensory representation. These data suggest that modulation of TH^VTA^ neurons can impact brain dynamics across many distributed anatomically distinct regions, even those that receive little to no direct TH^VTA^ input.

---

Midbrain dopaminergic neurons encode reward prediction errors^1-3^ and signal the incentive salience^4^ of sensory cues. During reward-seeking behavior, burst firing of these neurons results in phasic dopamine release^5-7^ in cortical and limbic terminal fields such as the medial prefrontal cortex (mPFC) and nucleus accumbens (NAc), which in conjunction with other neurotransmitters act to modulate postsynaptic neuronal firing^8,9^ and promote changes in motivated behavioral output^10-13^. The direct consequences of dopamine signaling are largely restricted to brain regions that contain appreciable presynaptic fibers that release dopamine, as well as postsynaptic dopamine receptors. However, dopaminergic signaling may also indirectly influence the activity in multiple brain regions, some of which may receive little to no direct dopamine input. By sculpting activity dynamics of neurons that are polysynaptically downstream, VTA DAergic neurons may thus modulate a much larger functional brain circuit. To test this hypothesis, we employed *in vivo* optogenetics coupled with fMRI^14-17^ to determine whether selective stimulation of TH^VTA^ neurons produced changes in fMRI signals and altered activity within multiple, anatomically distinct brain regions.

To selectively stimulate TH^VTA^ neurons *in vivo*, we introduced the light-gated cation channel, channelrhodopsin-2 conjugated to enhanced yellow fluorescent protein (ChR2-eYFP) selectively into VTA neurons in transgenic rats that express Cre recombinase under the control of the *tyrosine hydroxylase* (TH) gene promoter^18^ using established viral procedures^19^ (Fig. 1a). Because of a recent report of ectopic targeting of VTA neurons in TH-Cre mouse lines^20^, we performed a detailed quantitative assessment of viral targeting to TH+ neurons in the TH-Cre rat line across the anterior/posterior and medial/lateral subregions of the midbrain (Fig. 1a-c). We quantified the location of all neurons within anterior, middle, and posterior sections expressing only eYFP (eYFP+/TH-), TH (eYFP-/TH+), and neurons that expressed both (eYFP+/TH+) (n = 9 sections in n = 3 rats). Collapsed across the anterior-posterior axis containing the VTA, we observed highly specific viral transduction of TH+ neurons (97.4 +/-1.0%, n = 9 slices from n = 3 rats, Fig. 1d). Notably, few neurons that were eYFP+, but showed TH immunoreactivity that could not be resolved above background tended to reside in the anterior VTA or ventral to the VTA in the interpeduncular nucleus (Fig. 1c). In TH-Cre rats injected with AAV-DIO-ChR2-eYFP, we also observed substantial innervation of striatal subregions including the dorsal medial striatum (DMS) and the NAc (Fig. 1e,f). These results further validate the use of TH-Cre rats for targeted manipulation of TH^VTA^ neurons, with minimal targeting to TH-neurons in and around the VTA.

**Figure 1:**
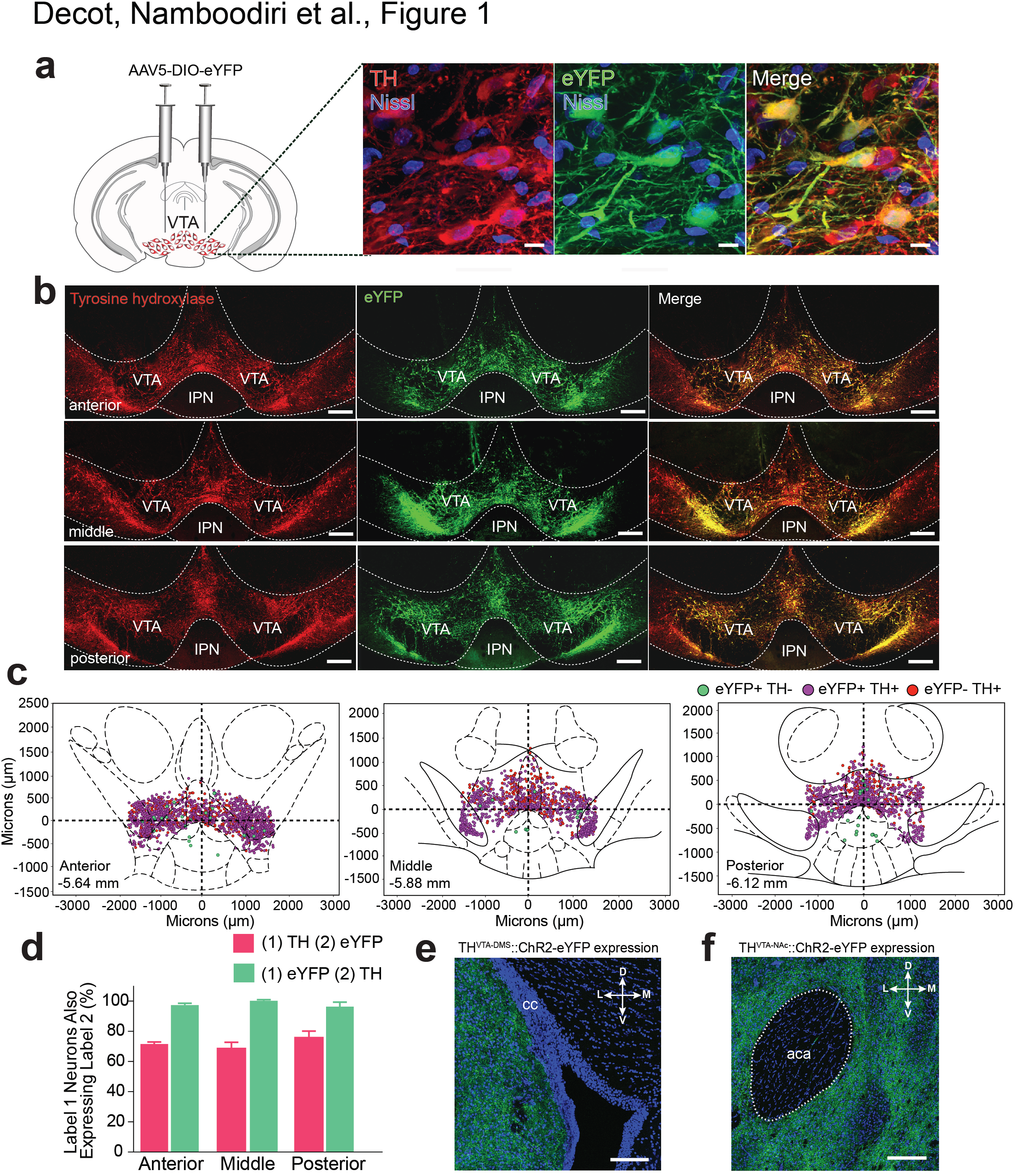
Viral targeting of VTA dopaminergic neurons in TH-cre rats. **a.** 63x and **b.** 20x confocal images depicting eYFP expression (green) following injection of a Cre-inducible virus into the VTA in TH-cre rats. Merged images reveal co-localization of eYFP and TH; red = TH; blue = Nissl stain; VTA: Ventral Tegmental Area, IPN: Interpeduncular Nucleus. **c,d.** Quantification of the location of all neurons within anterior, middle, and posterior slices of the VTA expressing only eYFP (eYFP+/TH-), TH (eYFP-/TH+), and neurons that expressed both (eYFP+/TH+) (*n* = 9 sections in *n* = 3 rats). Collapsed across the anterior-posterior axis, 71.9 ± 2.1 % of TH+ neurons within the VTA also expressed eYFP and 97.4 ± 1.0% of eYFP positive cells also expressed TH. Error bars represent SEM. e,f. Confocal images showing expression of ChR2-eYFP fibers in downstream target regions of the VTA including the dorsomedial striatum (DMS) and nucleus accumbens (NAc) of a *TH*::^VTA^::ChR2 rat. cc: corpus callosum, aca: anterior commissure. Scale bars, 200 and 500 µm, respectively.

In separate rats, we expressed ChR2-eYFP or eYFP alone in TH^VTA^ neurons and implanted bilateral optical fibers directly above the VTA for light delivery^21^ (Extended Fig. 1). Following recovery from surgery and sufficient time for adequate ChR2-eYFP expression in TH^VTA^ neurons, sedated rats were interfaced with a custom-built surface coil, placed in a 9.4 Tesla small animal MRI scanner, and the implanted optical fibers were connected to 473 nm diode pumped solid state lasers located outside of the MRI room (Fig. 2a). Following acquisition of 12, coronal T_2_-weighted anatomical image sections from each animal (Extended Fig. 1b), CBV-weighted fMRI^22-25^ data, which indirectly measures neuronal activity by detecting changes in hemodynamic signals accompanying vascular responses, were acquired using single-shot echo planar imaging sequence at 1 s temporal resolution. During these scans TH^VTA^ neurons were optically stimulated at 10 - 40 Hz for 10 s (Extended Fig. 2a,c). With these parameters we measured CBV responses across 12 continuous coronal slices of 1 mm thickness each encompassing nearly the entire cerebrum (see methods). Using a standard generalized linear model (GLM) to compare each voxel’s activity trace to a predefined stimulation template or hemodynamic response function^14,26,27^ (Extended Fig. 2e), optogenetic stimulation of TH^VTA^ neurons produced pronounced CBV increases in striatal brain regions in a frequency dependent fashion relative to control animals that received laser-light delivery, but only expressed eYFP in TH^VTA^ neurons (Extended Fig. 2). Further, systemic administration of a dopamine D1-receptor (D1R) antagonist, SCH23390, significantly attenuated CBV signals caused by TH^VTA^ optogenetic stimulation (Extended Fig. 2b). Taken together, these data show that selective activation of TH^VTA^ neurons produces dopamine receptor dependent signaling that result in increased CBV signals in dopaminergic terminal fields.

**Figure 2:**
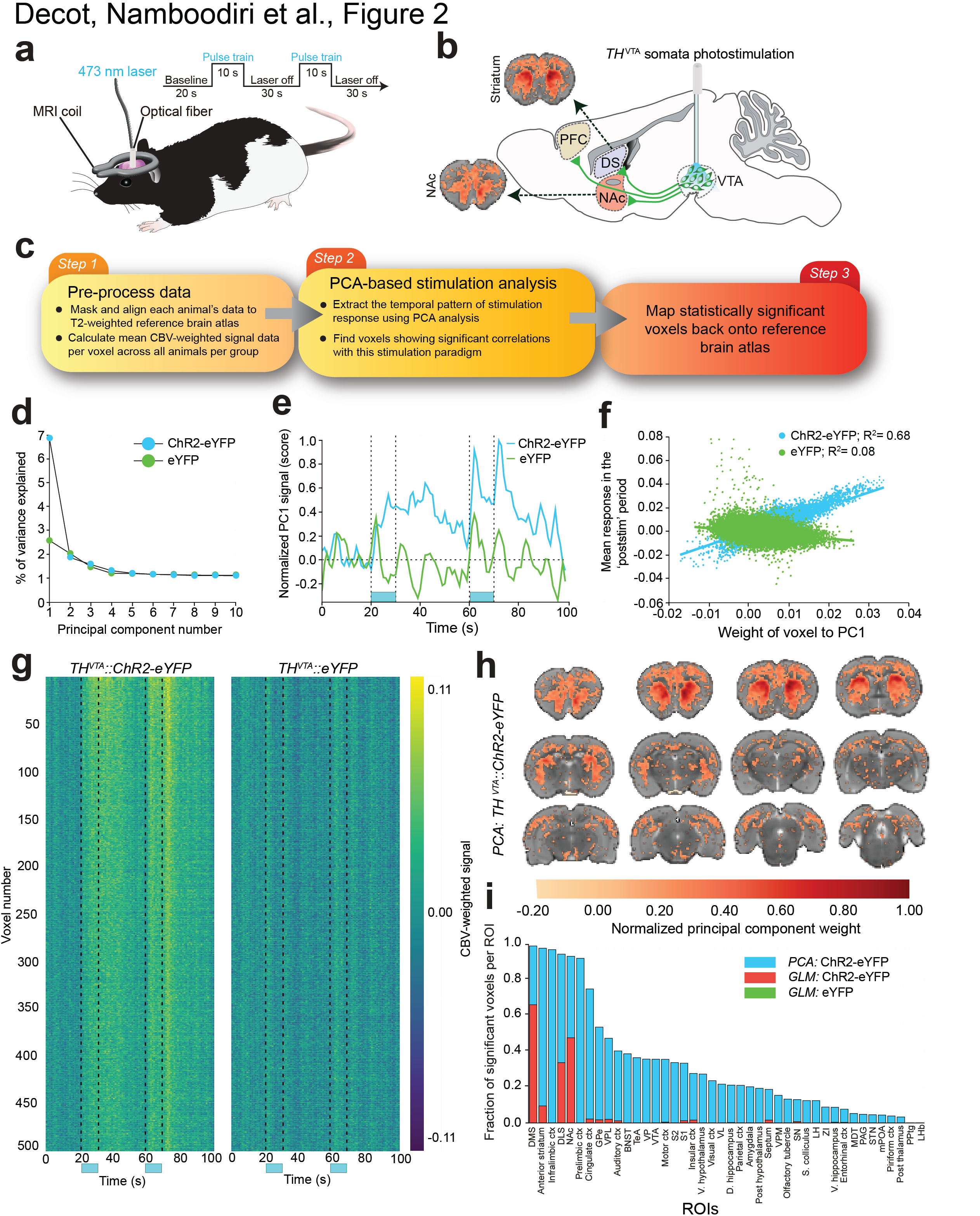
Whole brain analysis of the effect of *TH*^VTA^ neuron stimulation. **a-b.** Diagram of the experimental setup, **c.** Schematic of the analysis pipeline (see Methods), **d.** The percentage of variance captured by each principal component (PC) showing that the difference in variance explained between consecutive PCs is minimal after 2-3 PCs. **e.** The normalized trace (score) of the first PC for both experimental and control groups showing clear post-stimulation response only for the experimental group. The normalization was done such that the maximum score across both PCs is 1 (at 72s for the experimental group), **f.** A scatter plot showing the correlation between the weight of a voxel to the first PC and its mean post-stimulation response for both groups, **g.** The raw voxel traces for the top 500 voxels contributing to the first PC for both groups, **h.** The map showing the significant voxels contributing to the first PC vector for the experimental group, for which a significant post-stimulation response was identified (see Methods). There was no PC for the control group that represented any significant post-stimulation effect. PC weights were divided by the maximum PC weight for normalization, i. Individual ROIs and their fractional contribution (see Methods) to the PC representing the stimulation effect is shown in cyan. Red and green show the fraction of significant voxels that shows significant GLM coefficients for the experimental and control group, respectively.

Due to the lack of a well-defined dopamine-evoked hemodynamic response function and to mitigate the biases and assumptions imposed by traditional fMRI whole brain signal analysis, we developed an analytical pipeline by which all imaging voxels (35,182 per brain) and associated CBV weighted time series data were subjected to principal component analysis (PCA) factorization^28,29^ (Fig. 2c, Methods, **Supplementary Materials**). This resulted in a substantial reduction of the dimensionality of the spatial-temporal dataset from 35,182 down to 75 PCs that explained 80% of the variance of the data. Importantly, the first PC alone explained 7% of the variance of data collected from ChR2-eYFP rats. The percent variance of the data explained by each subsequent PCs reached an asymptote by 3-4 PCs (Fig. 2d).

The first PC plotted as a function of time represented the effect of TH^VTA^ optogenetic stimulation as it significantly increased from the baseline period (0 - 20 s) in each post-stimulation period (30 - 60 s, and 70-100 s) (t(48.0) = 11.69, p<0.001, Welch’s t-test for post-stimulation period 1 and t(38.7) = 7.15, p<0.001, Welch’s t-test for post-stimulation period 2, Fig. 2e), largely recapitulating the ROI analysis of striatal signals shown in Extended Fig 2a. In contrast, data collected from rats only expressing eYFP in TH^VTA^ neurons showed no significant post stimulation response. As the weight of a voxel to a given PC reflects the correlation between the voxel’s and the PC time series signals^30^, we hypothesized that a given voxel’s contribution to PC1 would highly correlate with the raw signal intensity for that voxel in the post-stimulation period exclusively in data collected from ChR2-eYFP rats. Supporting this, a highly significant positive Pearson’s correlation between these two variables was observed in the ChR2-eYFP group (R^2^ = 0.68), slope = 1.12, p < 0.001). In contrast, the eYFP group showed a slight, but significant negative correlation (R^2^ = 0.08, slope = −0.28, p < 0.001; Fig. 2f). We then ranked all voxels as a function of PC1 weight and plotted the time series for the top 500 voxels for both ChR2-eYFP and eYFP groups (Fig. 2g). Comparing these results to the top 500 voxels ranked by their mean post-stimulation response shows a dramatic enhancement of the temporal structure of the data (Extended Fig. 3b) further supporting our PCA based analysis approach.

We then mapped the voxels that significantly contributed to the PC representing the stimulation effect back to a T2-weighted reference brain atlas (Fig. 2h). This also showed a dramatic enhancement of brain wide activation patterns evoked by the stimulation compared to standard GLM based significant voxels (Extended Fig. 3c,d). In order to then place all significant PC1 weighted voxels back to defined brain regions we annotated all 12 T_2_-weighed brain slices into 42 anatomical subregions based on two widely used rodent brain atlases^31,32^ taking into account the spatial resolution of fMRI (Extended Fig. 4). Extracted significant voxels (based on PCA and GLM) mapped to numerous diverse brain regions (Fig. 2i). All GLM extracted voxels largely fell into ROIs within striatal subregions and frontal cortices as predicted based on innervation patterns of TH^VTA^ neurons^33^ (Fig. 1e,f). However, significant PCA extracted voxels represented a substantially larger fraction of each ROI within these subregions, further highlighting the enhanced sensitivity of this method. Furthermore, many additional brain regions were identified with a high fraction of significant voxels including basal ganglia, basal forebrain, and sensory cortical regions (Fig. 2i). Collectively, this analytical approach revealed that optogenetic modulation of TH^VTA^ neurons produces dispersed effects not only in brain regions that directly receive TH^VTA^ input, but also in numerous accessory regions not typically associated with direct VTA DAergic modulation.

In pathological conditions, such as addiction and schizophrenia, aberrant dopaminergic activity may act to over-enhance the salience of particular intrinsic and extrinsic stimuli^4,34^. Non-specific stimulation of the VTA can also alter cortical plasticity^8,35-37^, however it is unclear whether explicit pairing of TH^VTA^ activity with a discrete sensory stimulus alters the brain-wide sensory representation of that stimulus. Thus, in a new cohort of rats we tested whether somatosensory representations measured with fMRI was affected by coincidental and selective TH^VTA^ activity (Extended Fig. 5). We first measured changes in CBV signals in response to a range of forepaw stimulation frequencies. We then paired a single forepaw stimulation frequency (9 Hz) with 30 Hz optogenetic stimulation of TH^VTA^ neurons 20 times (one pairing every 70 s). Following the pairing protocol, we re-assessed changes in CBV signals in response to all forepaw stimuli frequencies (Fig. 3a). Forepaw stimulation produced timelocked CBV increases in the contralateral somatosensory cortex ROI in a frequency dependent fashion. Following pairing of 9 Hz forepaw stimulation with TH^VTA^ activity, there was a selective enhancement in CBV signals in response to 9 Hz subsequent forepaw stimulation compared to other non-paired frequencies and data from control animals that received laser light delivery, but did not express ChR2 in TH^VTA^ neurons (Fig. 3b-e, Extended Fig. 6).

**Figure 3:**
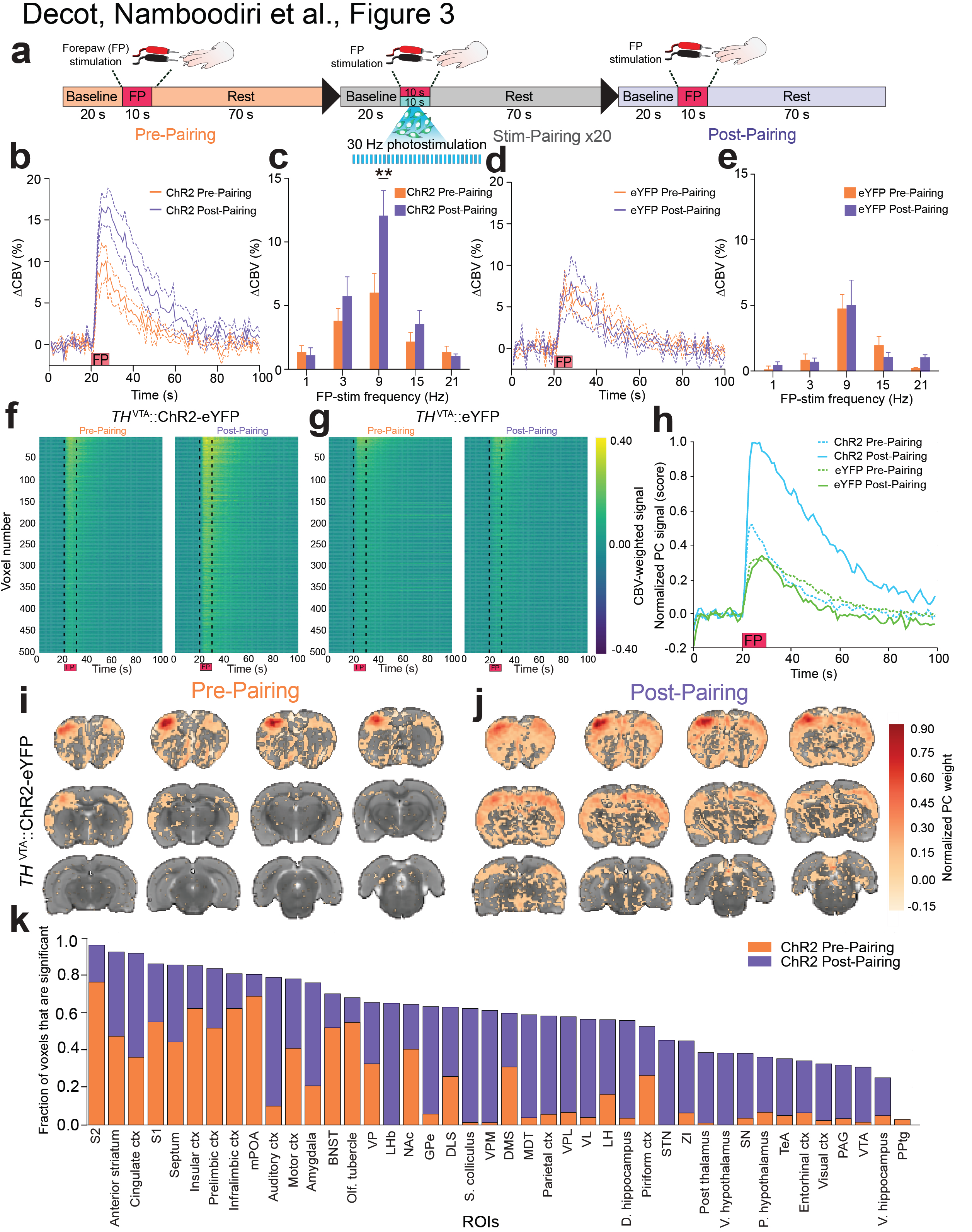
Whole brain analysis of forepaw stimulation pre and post pairing with *TH*^VTA^ neuron stimulation. **a.** Experimental timing diagram, **b.** Following pairing of VTA dopaminergic activity with 9 Hz FP stimulation there was a significant increased somatosensory cortex CBV timecourse signals (group main effect: *F*_1,99_ = 183.7, *P* < 0.0001, *n* = 7 rats per group), **c.** Pairing VTA dopaminergic activity with 9 Hz FP stimulation selectively enhanced somatosensory CBV signals at 9 Hz compared to other tested frequencies (pairing main effect: *F*_1,50_ = 5.58, *P* = 0.0219, *n* = 6-7 rats per group, double asterisks denote significant (p < 0.01) post-hoc tests between before and after pairing at each frequency), d. Following pairing of VTA laser light delivery with 9 Hz FP in rats only expressing eYFP did significantly change CBV responses in somatosensory cortex although the postpairing effect was slightly lower than prepairing (group main effect: *F*_1,99_ = 3.976, *P* = 0.0465, *n* = 5 rats per group), **e.** Pairing VTA laser light delivery with 9 Hz FP stimulation in rats only expressing eYFP did not alter somatosensory CBV frequency tuning signals (pairing main effect: *F*_1,50_ = 0.02, *P* = 0.8785, *n* = 5 rats). **f, g**. Raw voxel traces pre and post pairing around the forepaw stimulation (FP) for experimental and control groups, respectively. Post-pairing FP shows an enhancement of responses only for the experimental group. The voxels are sorted according to their pre-pairing FP response. The same row represents the same voxels across pre and post-pairing for both groups, **h.** PC traces (scores) for all conditions, **i, j.** The PC vectors capturing the FP response for pre and post pairing in the experimental group, showing a dramatic enhancement of voxels that significantly contribute to the PC representing the FP response, **k.** Individual ROIs showing fractional contribution to the PCs representing FP response.

To assess whole brain activity patterns before and after forepaw-TH^VTA^ pairing, we applied the PCA pipeline described above (Fig. 2c) to this dataset. PC1 signal intensity evoked by forepaw stimulation was significantly enhanced following pairing in the ChR2-eYFP group ChR2: (t(18) = 19.484, p < 0.001) but not to eYFP controls (t(18) = 0.660, p = 0.52 two-tailed t-test of the mean response in the stimulation period, Fig. 3f-h, Extended Fig. 7). Significant voxel weights to the PC1 vector remapped back to the reference atlas revealed a striking enhancement of the fMRI measured sensory representation of forepaw stimulation induced by the pairing protocol (Fig. 3i-k). To determine whether repeated TH^VTA^ forepaw stimulation pairing gradually enhanced the representation as a function of each pairing, we also calculated the PC fMRI signal for each TH^VTA^ forepaw pairing (20 total). Normalized to the maximum PC signal evoked by forepaw stimulation signals after the pairing, we observed that even the first and all subsequent TH^VTA^ forepaw stimulation pairings produced an increase in the PC signal in ChR2-eYFP rats relative to eYFP controls (Fig. 4a). This is further shown by plotting the ratio of each evoked response per pairing between ChR2-eYFP and eYFP groups (Fig. 4b). For either group, normalizing the PC signal to the respective pre-pairing forepaw response revealed a gradual habituation of the signal with increasing number of pairings (Fig. 4c). Collectively, these data suggest that forepaw stimulation paired with dopaminergic activity enhances the PC-fMRI signal gain during pairing as well as the subsequent brain wide representation of sensory stimuli.

**Figure 4:**
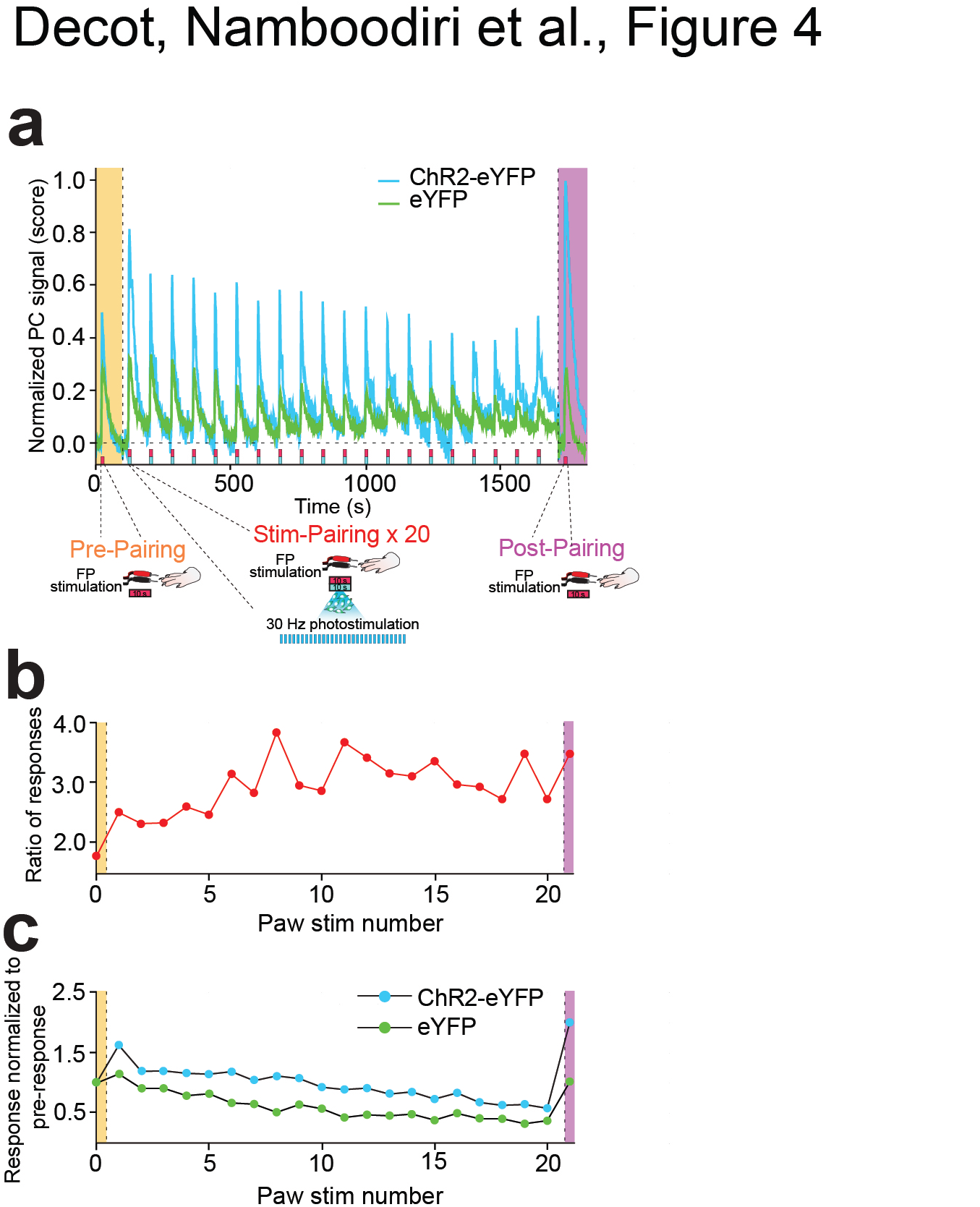
Evolution of forepaw stimulation pairing response over the course of pairings. **a.** The PCs identified (see Methods) as the ones representing forepaw stimulation response for both groups. The plots show pre-pairing and post-pairing forepaw stimulation responses, along with 20 responses during the pairing, **b.** The ratio of forepaw stimulation response (difference between the peak response during stimulation and the mean response during a 10 s baseline period prior to stimulation) for each stimulation between the ChR2-eYFP group and the eYFP group. There is a significant enhancement (positive linear regression) in this ratio as a function of the forepaw stimulation number (*slope* = 0.044, *P* = 0.005, *R*^2^ = 0.33). **c.** When the forepaw stimulation response for either group is normalized to the pre-pairing response for the same group, there is significant habituation (negative linear regression) with the number of stimulations during pairing for both groups (ChR2-eYFP: *slope* = −0.041, *R*^2^= 0.895; eYFP: *slope* = −0.034, *P* < 0.001, *R*^2^= 0.831). Importantly, the magnitude of habituation, as measured by the slope of response with respect to pairing number, is statistically indistinguishable for both groups (*F*_1,19_ = 2.10, *P*= 0.156, interaction term in ANCOVA).

While released dopamine acts to locally modulate neurocircuit function, our findings suggest that large-scale brain wide activity dynamics, measured across many distinct neuroanatomical regions, are also directly or indirectly regulated by activity of VTA dopaminergic neurons *in vivo*. In addition, dopaminergic activity, coincidental with a particular sensory stimulus, enhanced stimulus-specific neuronal representations brain wide. Previous work has largely focused on the timing at which dopaminergic activity occurs during ongoing behavior^1,2,5-7^ as well as cellular mechanisms by which dopamine signaling can alter the function of individual neurons^8,36,38,39^. Our results show that VTA dopaminergic activity has immediate global consequences, thereby potentially remodeling brain-wide neuronal activity patterns alone or in the presence of a sensory stimulus.

In order to detect and characterize brain-wide activity patterns measured with fMRI following TH^VTA^ optogenetic stimulation, we developed a PCA based approach. This allowed us to reduce the dimensionality of the data in the voxel space, isolate the PC temporal trace that represents the stimulation effect, and then map voxels that contribute significant weight to this PC back to anatomical locations in the brain. The identification of brain voxels that significantly contribute to the PC representing the stimulation effect is effectively similar to the standard GLM approach^27^ with notable distinctions. First, standard implementations of GLM require *a priori* specification of an expected stimulation effect. This is traditionally calculated using a hemodynamic response function. However, these functions vary widely across brain regions and species, and additionally are oftentimes derived from task-evoked activity, which is difficult to evaluate brain wide. Moreover, if multiple valid stimulation templates need to be compared (especially in the case with multiple regions of interest), power of the analysis can be dramatically compromised due to corrections for multiple comparisons. In contrast, our model free approach is entirely data driven and makes no assumptions of regional fMRI response specificity, thus making it highly applicable for analyzing brain wide changes in fMRI activity throughout the entire rat brain in response to timelocked stimulation. Another key benefit of our approach is apparent in the case where the stimulation of interest is not of a sensory region but is instead of a region that releases neuromodulators (e.g. VTA). In this case, modeling the expected stimulation effect as a standard sensory stimulation based template may be inappropriate as these models assume an exponential decay of the signal immediately following stimulation offset (Extended Fig. 2e).). However, extracting the stimulation response template using PCA (Fig. 2e) shows that the stimulation has a prolonged effect even after stimulation offset. Thus, in such cases, we advocate a data-driven approach to extract the stimulation response function. One caveat to our method is that a single PC representing the stimulation effect might also contain shared variation that is independent of the stimulation (for example intrinsic brain oscillations across regions). This is mitigated within our pipeline by averaging raw fMRI voxel across all animals in each group, since the shared temporal variability that remains in the data is likely only due to the stimulation. It is also possible that a single PC does not completely isolate the entire stimulation effect on brain wide activity. This in theory could be improved by additionally performing independent component analysis within the subspace formed by the top PCs. However, in the current dataset no poststimulation effects were observed outside of the first PC. Taken together, our brain wide fMRI analysis approach was designed to be easily implemented and sensitive enough to detect stimulation-induced changes in activity across the brain, while also imposing as few assumptions as possible.

By elucidating the brain wide activity pattern evoked by TH^VTA^ stimulation paired with a sensory stimulus, we show that sensory plasticity in this experiment is not limited to sensory cortices, but instead recruits numerous extrasensory brain regions. The dramatic sensory pattern emergence following TH^VTA^-forepaw pairing showcases the utility of optogenetic-fMRI experimentation (Fig. 3), as findings would have likely been unresolvable using methods that are focused on recording activity in one particular anatomical site. Importantly, our approach can easily be applied to examine the brain wide consequences of the modulation of any particular neurocircuit node. Our data also suggest that by pairing a sensory stimulus with activation of the TH^VTA^ neurons produces an immediate enhancement of the sensory representation (even on the first pairing) while not affecting the rate of habituation to the sensory stimulus (Fig. 4a-c). Our findings provide a foundation and roadmap for future studies that will examine the duration and/or permanence of the brain-wide sensory enhancement by TH^VTA^ stimulation as well as examine similar phenomena at the cellular level in a particular brain regions using large-scale single cell calcium imaging approaches.

A hallmark feature of both adaptive learning and addiction is the enhanced salience that sensory stimuli acquire when paired with natural rewards or drugs of abuse. Consistent with this, direct activation of VTA dopaminergic neurons promotes stimulus driven learning^3,12,18^, but can also result in addiction-like phenotypes in animals^40^. While the mechanistic explanation for these phenomena have largely focused either on VTA dopaminergic neurons or downstream targets such as the NAc, our data suggests that adaptive and/or maladaptive functioning of the dopamine system not only biases activity within mesolimbic reward circuits, but also promotes brain-wide changes in network dynamics, to generate behavioral output.

## Acknowledgements

We thank members of Stuber and Shih labs for discussion. We thank the UNC vector core for viral packaging, and the UNC Neuroscience Center Microscopy Core (P30 NS045892). This study was supported by The Brain and Behavior Research Foundation, The Foundation of Hope, The Klarman Family Foundation, and the National Institute on Drug Abuse (DA032750, DA038168) (G.D.S.), and UNC Neurology-BRIC startup funds, University Research Council, and SeeCure LLC (Y.Y.I.S.). H.K.D. was supported by NS007431 and UNC Graduate Training Program in Translational Medicine supported by HHMI. J.H.J. was supported by MH104013. M.D. was supported by a Human Frontier Science Program Cross-Disciplinary Fellowship.

### Author contributions

H.K.D., Y.Y.I.S., and G.D.S. designed the experiments. H.K.D., J.H.J., J.A.M., P.A.K., Y.C.J.K., and M.D. collected data. K.D. and I.B.W. provided critical reagents. V.M.K.N., H.K.D., W.G., S.L., Y.Y.I.S., and G.D.S. analyzed data. G.D.S., H.K.D., and V.M.K.N. wrote the manuscript with input from all authors.

## Methods

### Experimental subjects and stereotactic surgery

All procedures were conducted in accordance with the Guide for the Care and Use of Laboratory Animals, as adopted by the National Institutes of Health, and with approval of the Institutional Animal Care and Use Committee at the University of North Carolina (UNC). Adult (400-450 g) male tyrosine hydroxylase (TH)-ires-cre Long Evans rats were group housed until surgery and were maintained on a 12-h light cycle (lights off at 19:00) with mild food restriction to maintain ~90% body weight during the duration of the study. To target dopamine (DA) neurons within the midbrain, TH-ires-cre rats were endotracheally intubated and ventilated using a small animal ventilator (CWE Inc., SAR-830/PA, Armore, PA) with ~1.5% isoflurane in medical air prior to being placed into a stereotactic frame (Model 962, Kopf Instruments, Tujunga, CA). For all experiments, rats were microinjected with quadruple injections of 1 μl of purified and concentrated adeno-associated virus (~10^12^ infections units per ml, packaged by the UNC Vector Core Facility) into the ventral tegmental area (VTA) using the following coordinates (in mm from bregma): −5.4 and −6.2 anterior/posterior, ±0.7 medial/lateral, −8.4 and −7.4 dorsal/ventral. VTA dopaminergic neurons were transduced with an AAV5 carrying a cre-inducible expression cassette encoding channelrhodopsin-2 (ChR2) fused to an enhanced yellow fluorescent protein (eYFP) under the control of the EF1α promoter (AAV5-DIO-ChR2-eYFP; TH^VTA^::ChR2 rats) or only eYFP (AAV5-DIO-eYFP; TH^VTA^::control rats). 200 μm multimode chronic optical fibers were stereotactically implanted bilaterally at a 10° angle directly above the VTA using the following stereotactic coordinates: −5.8 mm to bregma, ±2.14 mm lateral to midline, and −7.8 mm ventral to the skull surface. The time from virus injection to the start of the MRI experiments was 5–6 weeks for all subjects.

### fMRI procedures

MRI was performed using a 9.4 Tesla Bruker BioSpec system with a BGA-9S gradient insert (Bruker Corp., Billerica, MA) at the UNC Biomedical Research Imaging Center (BRIC). On the day of MRI experiments, each rat was endotracheally intubated and ventilated with ~1.5% isoflurane in medical air. The ventilation rate and volume were adjusted via a capnometer (Surgivet v9004, Smith Medical, Waukesha, WI) to maintain end-tidal CO_2_ (EtCO_2_) within a range of 2.9±0.3%. Non-invasive EtCO_2_ values were previously calibrated against invasive blood-gas samplings under identical baseline conditions, resulting in an arterial pCO_2_ of 37.6± 4.7 mmHg^1^. Heart rate and oxygen saturation (SpO_2_) were continuously monitored by a non-invasive MouseOx Plus System with MR-compatible sensors (STARR Life Science Corp., Oakmont PA) and maintained within normal ranges (350-400 bpm and above 96%, respectively). Rectal temperature was maintained at 37±0.5°C with a warm-water circulating pad. Black tape was placed over the eyes and a masking light was directed into the face of each rat to minimize visual stimulation during laser light delivery.

A home-made surface coil with an internal diameter of 1.6 cm placed directly over the head was used as a RF transceiver. Magnetic field homogeneity was optimized using standard FASTMAP shimming with first order shims on an isotropic voxel of 7x7x7 mm encompassing the imaging slices. A RARE T_2_-weighted image was taken in the mid-sagittal plane to localize the anatomical position by identifying the anterior commissure at 0.36 mm posterior to bregma. To ensure anatomical consistency of the imaging data, twelve coronal slices were acquired using T_2_-weighted imaging, with the 4^th^ slice from the anterior direction aligned with the anterior commissure. A RARE sequence (spectral width=47 kHz, TR/TE=2500/33 ms, FOV=2.56x2.56 cm, slice thickness=1 mm, matrix=256x256, RARE factor=8, and number of averages=8) was used to confirm optical fiber position with reference to the cortical surface, midline, and the anterior commissure. For functional scans, single shot gradient echo-EPI sequence (spectral width=300 kHz, TR/TE=1000/8.1 ms, FOV=2.56x2.56 cm2, slice thickness= 1 mm, matrix=80x80, providing temporal resolution=1 s) was used.

During fMRI, dexmedetomidine (0.1 mg/kg/hr) and pancuronium bromide (1.0 mg/kg/hr) were infused intraperitoneally and isoflurane was lowered to 0.5% at 30 min after the infusion began^2^. Cerebral blood volume (CBV)-weighted fMRI^3^ was achieved by injecting Feraheme (AMAG Pharmaceuticals, Lexington, MA) at a dose of 30 mg Fe/kg via a tail vein catheter. Optical fibers were coupled via 3-m patch cables to solid-state lasers (473 nm wavelength) located outside of the MRI room delivering ~10 mW light to each hemisphere of the VTA. Evoked fMRI scans were acquired for 100 s during which photostimulation was applied in a 20-s OFF, 10-s ON, 30-s OFF, 10-s ON, 30-s OFF pattern. Subjects underwent two to five repeated trials at each photostimulation frequency (10, 20, 30 and 40 Hz at 5 ms pulse width) in a pseudo-random manner. Dopamine D1 receptor antagonist, SCH23390 (0.6 mg/kg) (Sigma) was injected intravenously to explore its effects on stimulus-evoked CBV changes observed in TH^VTA^::ChR2 rats.

For paired forepaw and optogenetic stimulation experiments, electrical forepaw stimulation was applied using two needle electrodes inserted under the skin of the right forepaw of each subject. The stimulation was applied in a 20-s OFF, 10-s ON, 70-s OFF pattern. To measure the frequency dependence of the stimulus-evoked activation, subjects underwent two to five repeated trials at the following frequencies: 1, 3, 9, 15, and 21 Hz with a fixed pulse width of 0.3 ms and a current of 1.5 - 3.0 mA (pre-pairing). Then, a repeated pairing of forepaw electrical stimulation at 9 Hz with 30 Hz optogenetic stimulation of VTA DA neurons was performed with an initial 20-s OFF, followed by 20 blocks of 10-s ON and 70-s OFF. Finally, changes in CBV signals and frequency dependence in response to all forepaw stimuli frequencies were re-assessed (post-pairing).

### Data processing and analysis

fMRI data analyzed using a generalized linear model (GLM) were processed using Matlab (Math-Works, Natick, MA) and Statistical Parametric Mapping (SPM) codes, with the pipeline similar to our previous publications^4-7^. Automatic co-registration were applied to realign time-series data within subjects to correct subtle drift of EPI images and then again across subjects to provide group-based fMRI maps. Data were analyzed using the Analysis of Functional Neuroimages (AFNI) with the framework of GLM^8-10^, with false discovery rate (FDR) correction to adjust for the multiple comparisons of fMRI maps (p<0.05). The stimulation paradigm was convoluted with response model of Feraheme^11^ injection in AFNI package in order to get an ideal model of hemodynamic response (Extended Fig. 2e). The stimulus-evoked ΔR_2_* value was calculated as follows after Feraheme injection: 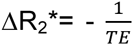 In (S_stim_/S_o_), where S_stim_ and S_o_ are the MR signal intensities before stimulation and before Feraheme injection, respectively. The effect of optogenetic and forepaw stimulation frequency on ΔCBV change was calculated by averaging CBV values from the first 20 s time points during each stimulation epoch. Regions of interest (ROIs) were placed on the dorsal striatum to extract fMRI time-course data after the images were co-registered. For pairing experiments, ROI was placed on the contralateral somatosensory cortex (S1).

PCA analysis pipeline for data analyzed utilizing this method within this manuscript was written in python 2.7 using numpy and scikitlearn packages. All analysis code is available in well-documented iPython notebooks (**supplementary materials**).

### Histology, immunohistochemistry and microscopy

Rats were deeply anaesthetized with pentobarbital, and transcardially perfused with phosphate buffered saline (PBS) followed by 4% (weight/volume) paraformaldehyde in PBS. Brains were post-fixed in 4% paraformaldehyde for 24 hr and transferred to 30% sucrose in ddH2O for 48 hr. 40 μm brain sections were collected and subjected to immunohistochemical staining for neuronal cell bodies (NeuroTrace Invitrogen; 435 nm excitation/455 nm emission and/or tyrosine hydroxylase) (Pel Freeze; made in sheep, 1:500). Z-stack and tiled images of mounted brain sections were visualized using Zeiss LSM 710 confocal microscope with a 20x or 63x objective.

---

**Extended Data Figure 1:**
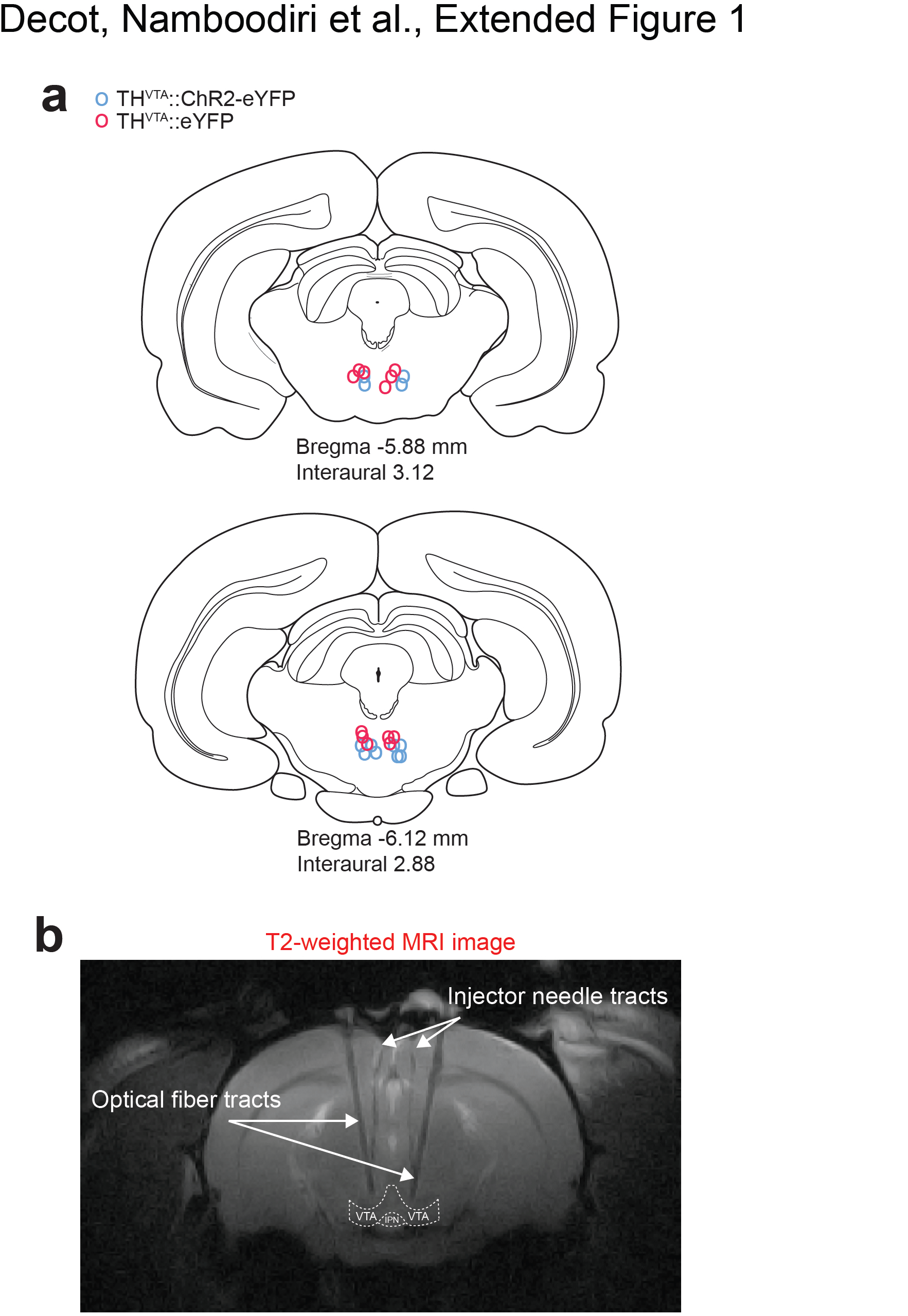
Optical fiber placements in the VTA of TH-cre rats. **a.** Location of bilateral optical fibers within the VTA of TH^VTA^::ChR2-eYFP and TH^VTA^::eYFP rats based on T2-weighted image examination and histological verification of 40-μm brain slices following experiments. **b.** RARE T2-weighted sequences were used to confirm optical fiber placements.

**Extended Data Figure 2:**
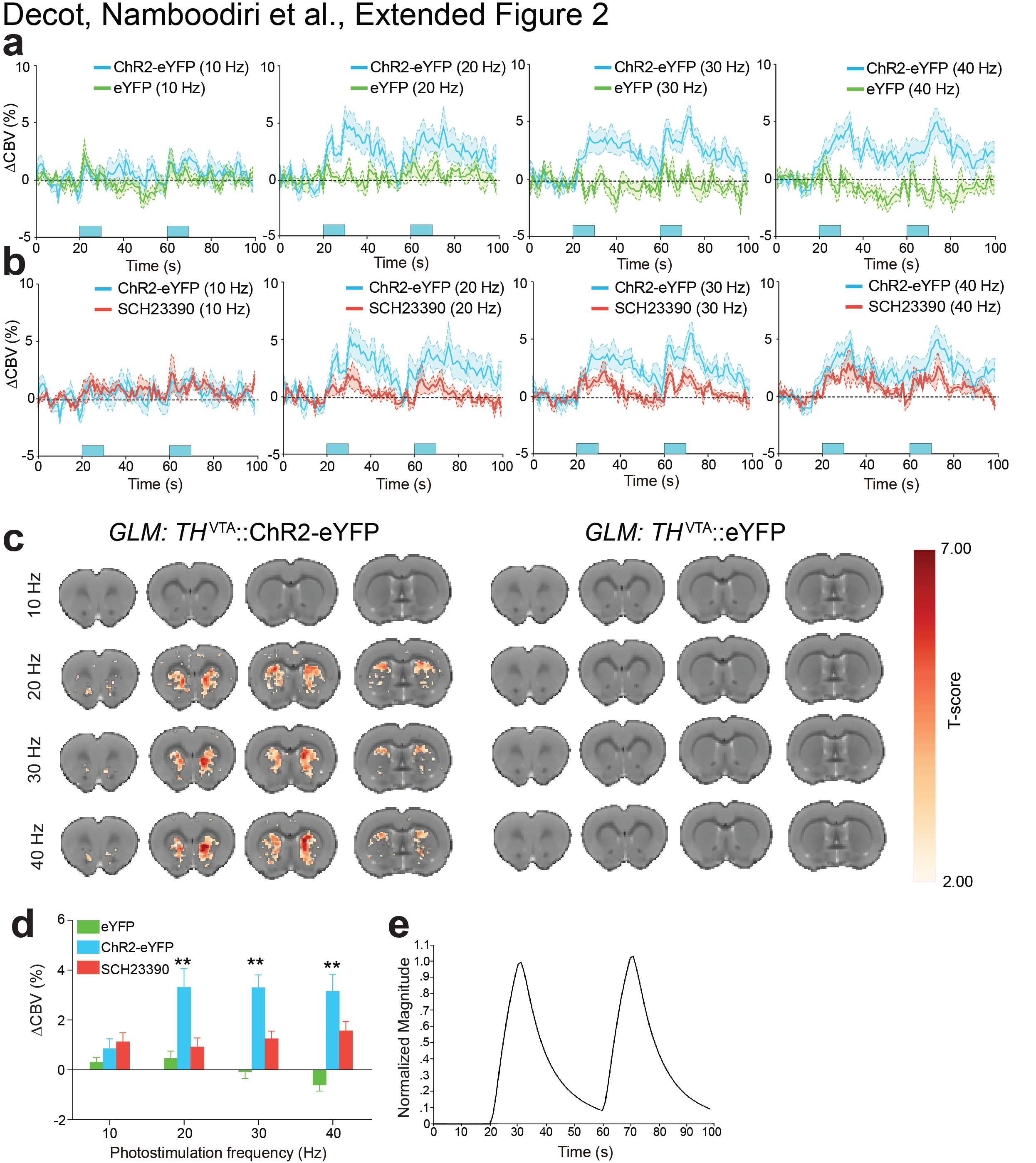
Activation of VTA dopaminergic neurons increases CBV signals in striatal target regions. **a.** Activation of VTA dopaminergic neurons increases striatal CBV signals compared to control rats, 20 Hz (group main effect: *F*_1,99_ = 204.6, *P* < 0.0001, *n* = 6 rats per group), 30 Hz (group main effect: *F*_1,99_ = 479.0, *P* < 0.0001, *n* = 6 rats per group) and 40 Hz (group main effect: *F*_1,99_ = 506.4, *P* < 0.0001, *n* = 6 rats per group). **b.** Intravenous administration of SCH23390 significantly attenuates optically-evoked changes in striatal CBV timecourse signals at 20 and 40 Hz, 10 Hz (group main effect: *F*_1,99_ = 1.874, *P* =0.1714, *n* = 5 - 6 rats per group), 20 Hz (group main effect: *F*_1,99_ = 191.2, *P* < 0.0001, *n* = 5 - 6 rats per group), 30 Hz (group main effect: *F*_1,99_ = 226.7, *P* < 0.0001, *n* = 5 - 6 rats per group), and 40 Hz (group main effect: *F*_1,99_ = 92.76, *P* < 0.0001, *n* = 5 - 6 rats per group). **c.** Group-averaged CBV activation maps following optogenetic stimulation of VTA dopaminergic neurons. **d.** Activation of VTA dopaminergic neurons increases striatal CBV signals in a frequency dependent fashion compared to SCH23390 and eYFP groups (interaction main effect: *F*_6,124_ = 3.573, *P* = 0.0027, *n* = 5 - 6 rats per group, double asterisks denote significant (p < 0.001) post-hoc effects of ChR2-eYFP vs. eYFP at each frequency). For all average data figures, mean and SEM are shown. **e.** Stimulation template assumed for the GLM analysis (Leite, F.P. et al., Repeated fMRI using iron oxide contrast agent in awake, behaving macaques at 3 Tesla. *Neuroimage* 16: 283-294 (2002).).

**Extended Data Figure 3:**
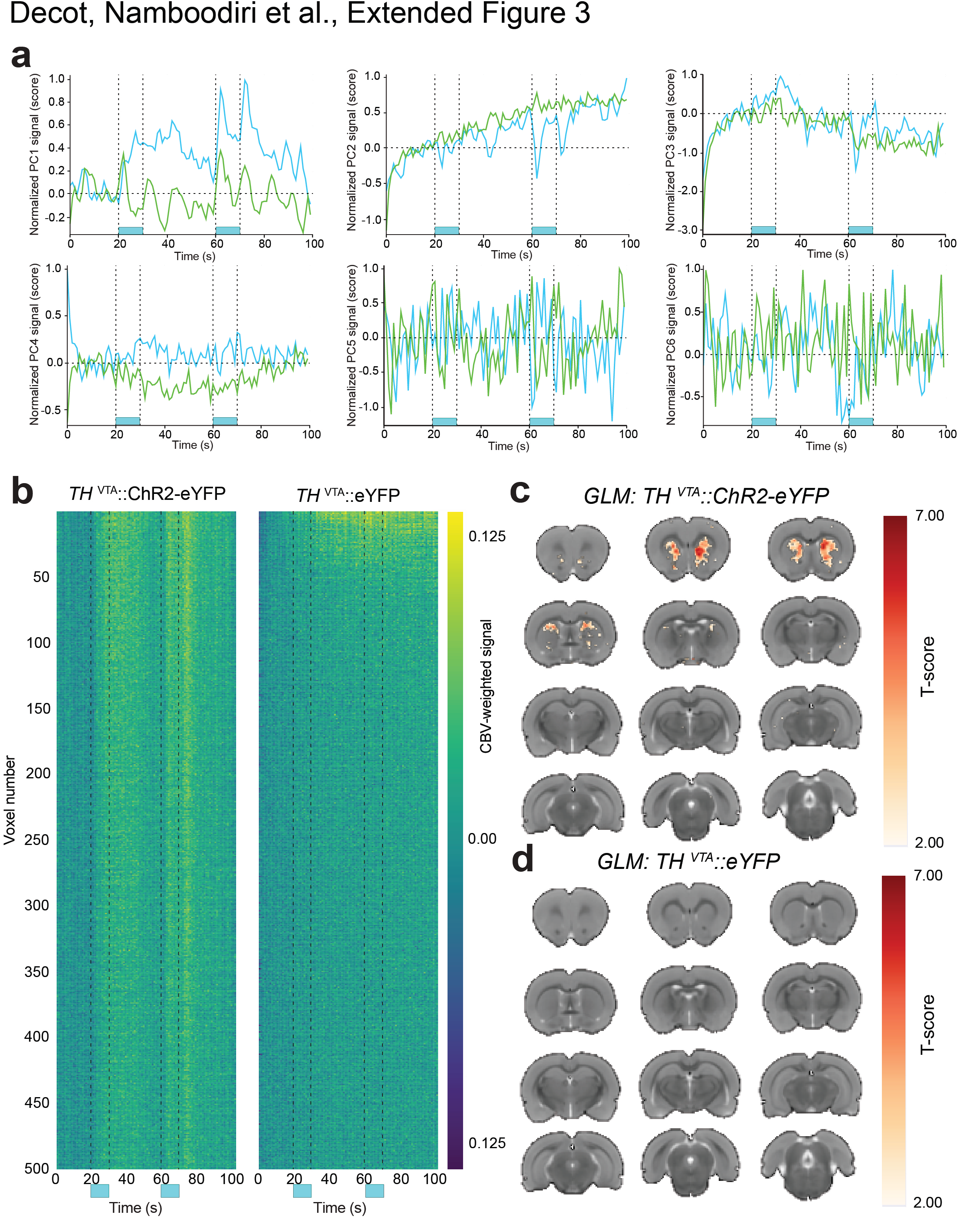
Additional results for the TH^VTA^ stimulation experiment. **a.** PC traces from the top 6 PCs for both groups. PCs beyond PC4 lack temporal structure. The first PC clearly represents stimulation effect, including a significant post-stimulation effect. The second PC largely represents a slow drift in the signal throughout the experimental session. PC4 for both groups are actually flipped in sign with respect to each other. Our sign standardization (see Methods) procedure enforced the change in signal at the first stimulation onset to be positive. This change is in opposite directions for the ChR2-eYFP and eYFP groups, which is why the traces are flipped. **b.** Raw fMRI signal from the top 500 voxels that showed the highest mean post-stimulation response. When compared to Figure 3e, it can be seen that ranking by PC weights reveals substantially more temporal structure in the responses. **c,d.** Activation maps due to optogenetic stimulation as assessed by a GLM of the fMRI response. The GLM coefficients are shown only for significant voxels following Benjamini-Hochberg false discovery rate correction.

**Extended Data Figure 4:**
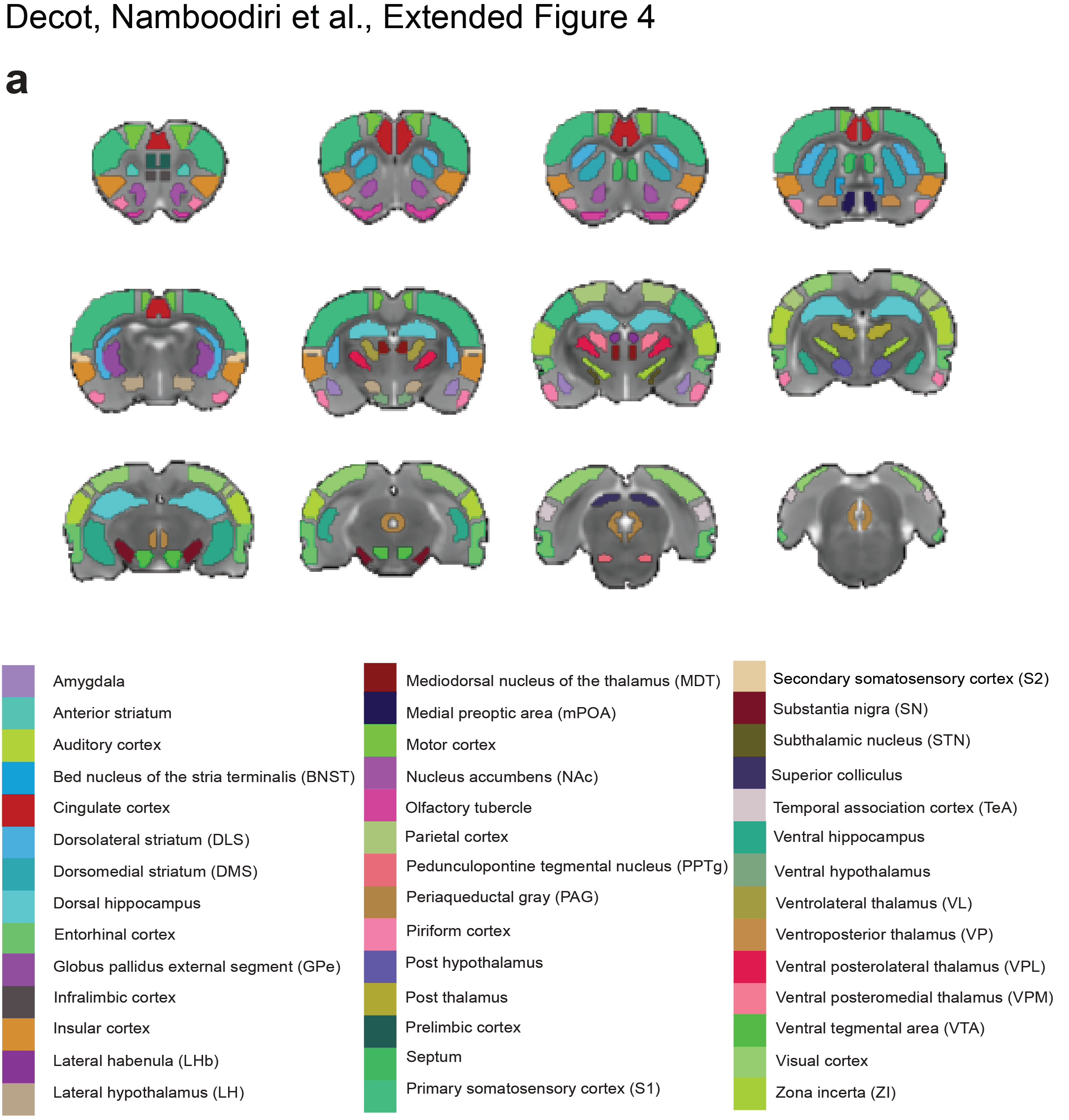
Map of defined regions of interest. 44 anatomical subregions are shown across the 12 slices of the brain.

**Extended Data Figure 5:**
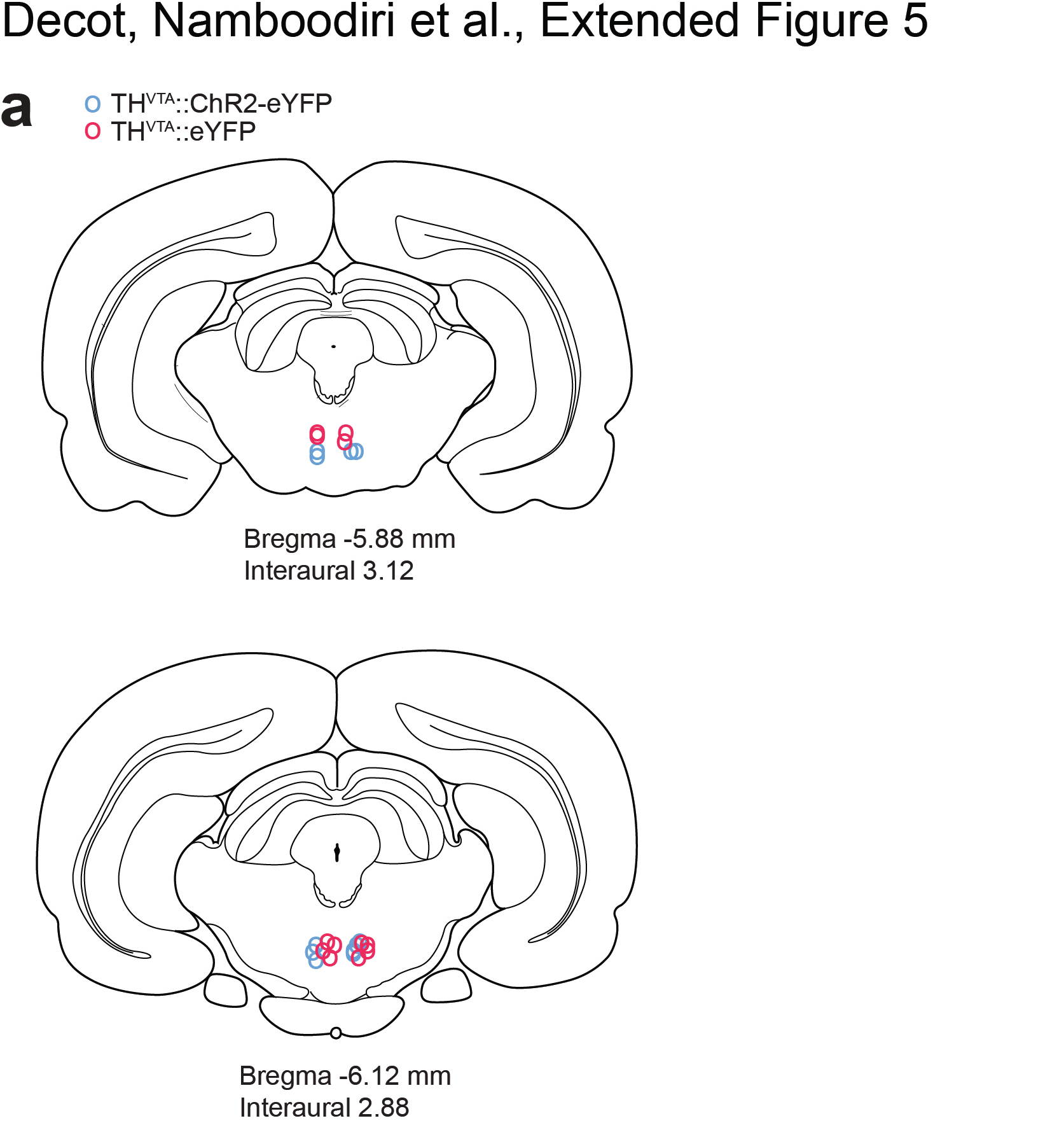
Optical fiber placements in the VTA of TH-cre rats for the pairing experiments. **a.** Location of bilateral optical fibers within the VTA of TH^VTA^::ChR2-eYFP and TH^VTA^::eYFP rats based on T2-weighted image examination and histological verification of 40-μm brain slices following experiments.

**Extended Data Figure 6:**
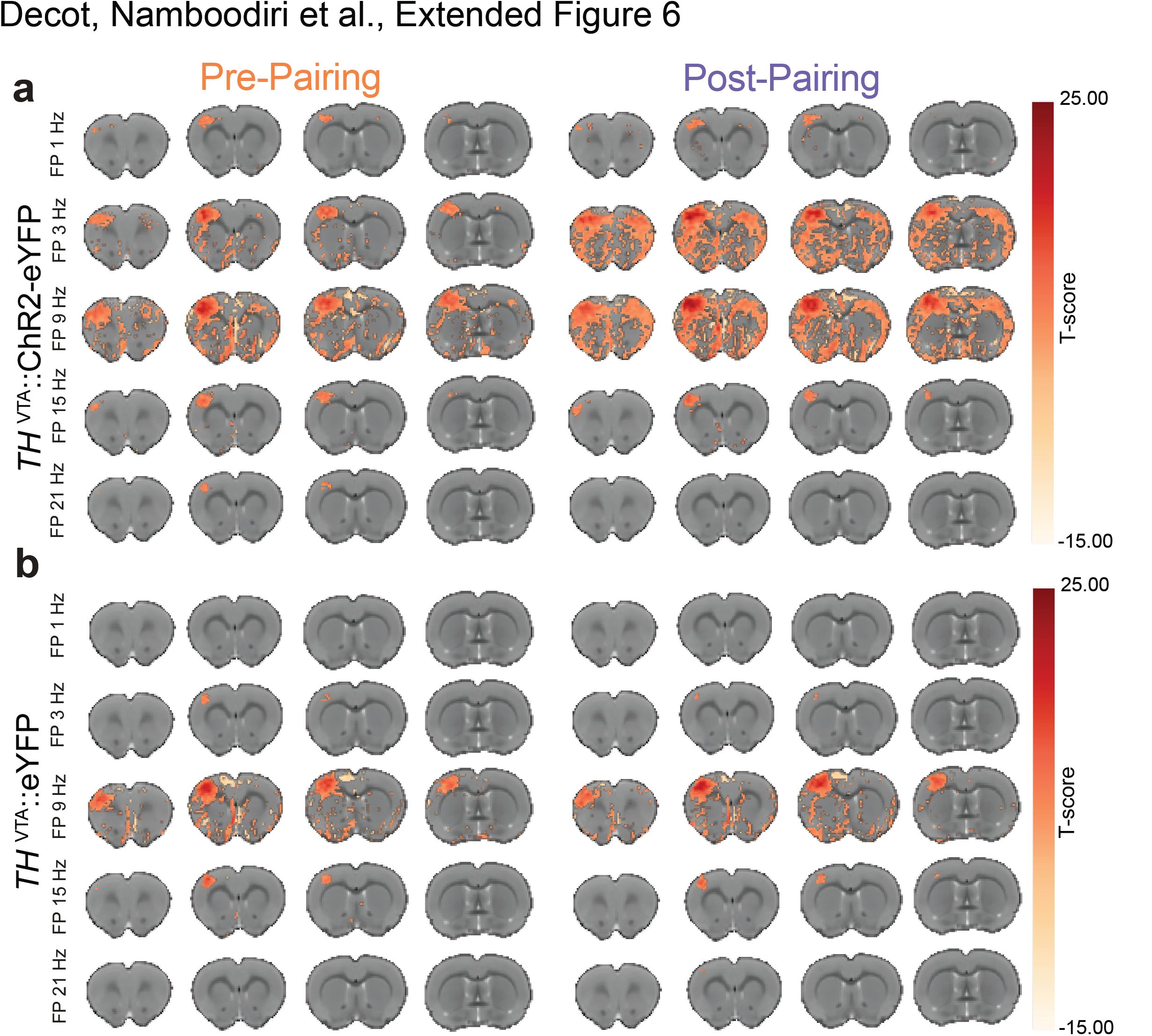
Pairing VTA dopamine neuronal activity with somatosensory stimuli enhances the neuronal representation of the sensory stimulus. **a,b.** Group averaged CBV activation maps in response to forepaw (FP) stimulation before and after pairing 30 Hz optogenetic stimulation of VTA dopaminergic neurons with 9 Hz FP stimulation for TH^VTA^::ChR2-eYFP and control rats.

**Extended Data Figure 7:**
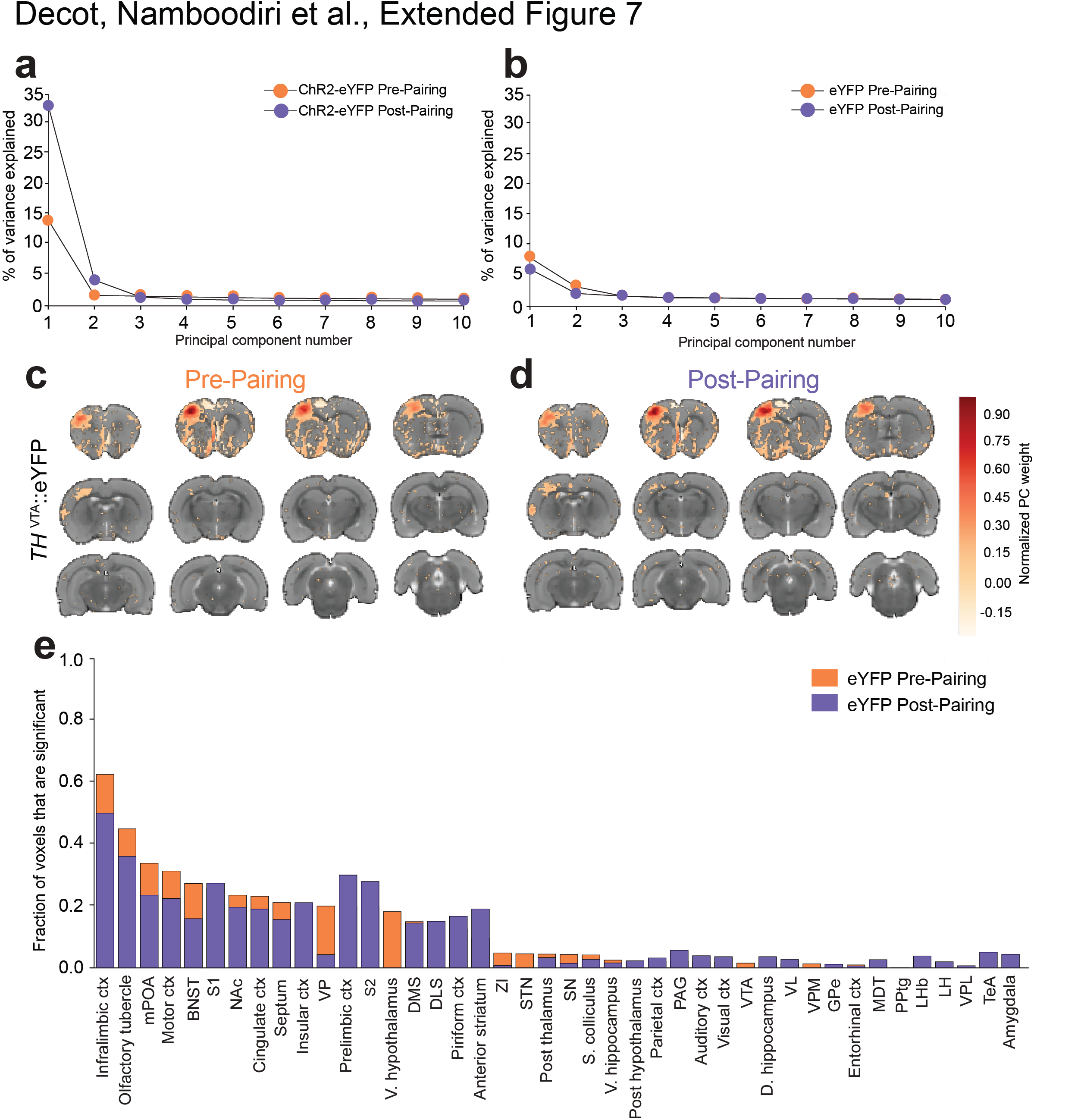
Additional results for the forepaw stimulation pre and post pairing with TH^VTA^ stimulation. **a, b.** Percentage variance explained per PC is plotted for the top 10 PCs for both groups pre and post pairing. The first PC represented the stimulation effect in all conditions. **c, d.** Significant voxels contributing to the PC vector representing stimulation effect are shown for the eYFP group pre and post pairing with TH^VTA^ stimulation. **e.** Ranked ROIs based on the fractional contribution of significant voxels to the PC representing stimulation effect for the eYFP group pre and post pairing.

## Analysis of fMRI data

This document details the PC-fMRI analysis pipeline used in Figs. 2, 3, **and** 4 in the manuscript. Additional ipython notebooks with documented code used for this pipeline can be found at https://github.com/stuberlab/<u>PC opto fMRI analysis</u>. Raw data used for this study can be downloaded via the associated ipython notebooks. The aim of the data analysis pipeline was to analyze the activity of the whole brain (as opposed to pre-defined ROIs) in relation to the various stimulation paradigms. To this end, we employed a Principal Components Analysis (PCA) based dimensionality reduction. We will first present the general outline of the pipeline before explaining the details of the different steps.

**Outline:**

1. Align and mask all animals’ data to the atlas.
2. For each animal, calculate the cerebral blood volume-weighted signal (adapted from^1^) for each voxel for each experimental run.
3. Calculate the mean data per voxel across all animals and runs.
4. Perform PCA on this mean dataset to reduce the dimensionality of the voxel space.
5. Select the principal component (PC) representing the stimulation response.
6. Calculate the voxels that significantly contribute to this PC (equivalent to a GLM that tests a linear relationship between the PC stimulation trace and each voxel’s trace).
7. Map these significant voxels back to pre-defined regions of interest (ROIs) and rank ROIs based on the fraction of voxels significantly contributing to the PC.

**Detailed description:**

1. We aligned the data of each animal to a reference T2 weighted atlas and then masked out the data outside of the reference atlas. This was done so as to line up the voxels across all animals.
2. In order to correct for the differences between animals introduced by differences in the injection of monocrystalline iron oxide nanoparticles (MION), we first normalized each animals’ fMRI signal by calculating the corresponding CBV-weighted signal as:

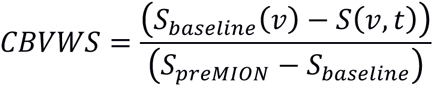

where *S*_*baseline*_(*v*), the baseline signal for voxel *v*, was calculated as the mean signal in the first 20s of the run for that voxel. *S*_*baseline*_in the denominator was calculated as the mean*S*_*baseline*_(*v*)across all voxels, and*S*_*preMION*_was calculated as the mean signal in the first 20s of the baseline scan prior to the injection of MION across all voxels. This analysis corrects for the average change in the signal introduced by the MION injection for each animal, before averaging the data from all animals together in the next step.
3. Prior to performing PCA, we averaged the traces per voxel across all animals and runs of an experiment. The primary purpose of this step was to ensure that only shared variation across animals (likely from stimulation) is present in the dataset on which PCA is performed. In this way, variability in the data extraneous to that introduced by stimulations would get averaged out. As a result, the extracted PC representing stimulation effect is likely going to have minimal contribution from noisy fluctuations in the signal.
4. Once the mean data for each group (CHR2 or EYFP) was calculated, we performed PCA on both of these datasets. In each case, the spatial dimensions representing the voxels were treated as the features and the temporal dimension treated as samples within this feature space. We also performed dimensionality reduction by limiting the number of principal components to that required in explaining 80% of the variance in the data. The PCA decomposition was done using scikit-learn in Python. Since the PCs are determined only up to a sign (180° rotation, for instance, does not change the PC), we performed a sign-standardization. This was done by requiring that the change in the signal at the moment of first stimulation is positive, i.e. if the change in the signal at the moment of first stimulation is negative, we multiplied the PC vector and trace (or score) by −1. Thus, the stimulation response in all PCs are positive; the caveat is that for PCs not representing stimulation effect, the change in the signal at this might be due to random fluctuations.
5. Having performed PCA, we then selected the PC that represented the stimulation effect. We defined this PC as the one that showed a significant difference in the mean signal during the post-stimulation period compared to the mean signal in the baseline period (using a Welch’s t-test). We also placed an additional constraint that the post-stimulation response trended towards the baseline. The latter criterion ensured that PCs showing a stable drift across the entire session were not considered as stimulation-induced. For the VTA stimulation experiment, each experimental run had two stimulations. Thus, in this case, we required that the PC representing the stimulation response satisfied the above criteria for both stimulations, i.e. the mean post-stimulation response was higher than the mean baseline response for both post-stimulation periods, and that the response during the post-stimulation period for both stimulations trended towards baseline. A caveat for this method is worth mentioning: it is possible that the stimulation effect is not *entirely* isolated within a single PC. In our case, it was clear looking at the traces of each PC that the post-stimulation effect is captured within the identified PC. If this was not the case, it would be appropriate to run a subsequent rotation of the axes within a subspace spanned by the first few PCs to isolate non-orthogonal components^2^. Subsequently, the result of this trace could be used as a stimulation model for a GLM approach. Since our analysis, on the other hand, was stopped at PCA, performing such a GLM is equivalent to identifying the significant voxels contributing to the PC of interest (see below).
6. We then calculated the voxels that significantly contributed to the PC representing the stimulation response calculated above. It is well known that the weight of a feature (voxel, in this case) to a PC can be represented as^3^.

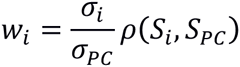

whereo*σ*_*i*_represents the standard deviation of the *i*^*th*^ voxel trace (*S*_*i*_),*σ*_*PC*_the standard deviation of the PC trace (*S*_*PC*_) and ρ, the Pearson correlation coefficient. Thus, the voxels that contribute significantly to the PC are the voxels that show significant Pearson’s correlation with the PC^3^. In other words, similar to General Linear Model (GLM) based approaches^4^, this method extracts out the voxels that show significant correlation with a stimulation response function. However, unlike GLM, we do not have to define a-priori models of stimulation effects using a putative hemodynamic response function (HRF). This is a really important advantage as it has been shown that HRFs are different depending on the species and region in the brain^5,6^. Crucially, the appropriate HRF for each voxel in the brain of rats is not known and may even be impossible to reliably specify. Thus, using PCA to instruct us the shape of the stimulation response is data-driven and mitigates biases introduced by the specifications of the GLM model.
7. Once the voxels showing significant correlation with the PC trace representing the stimulation effect are calculated as above, we mapped these voxels back to hand-drawn ROIs based on the reference atlas. Subsequently, we ranked the ROIs based on their fraction of voxels that significantly contributed to the PC representing the stimulation effect. Comparing ROIs based on the fraction of significant voxels has the benefit of normalizing for the size of the ROI. However, two caveats are worth mentioning: 1. This fraction ignores the magnitude of the response for each voxel, and, 2. ROIs with fewer voxels may have spuriously high or low fractions due to their small sizes. The raw data indicating the number of total and significant pixels for each ROI are uploaded as Supplementary Materials.

**Additional methods for Figure 4**

To analyze the effect of TH^VTA^ pairings with paw stimulation, we first concatenated the fMRI data from the pre-pairing epoch (averaged across animals and runs) with the data during pairing (averaged across animals) and the data from the post-pairing epoch (averaged across animals and runs). The above PCA pipeline was run on this combined dataset. To find the PCs representing paw stimulation response for both groups, we required two conditions to be met: a) there should be a positive stimulation response to every stimulation (22 in total), and b) the post-stimulation response should show a decreasing trend following every stimulation. The traces for the PCs identified thus are shown in Figure 4a. As these traces showed variable magnitudes prior to each stimulation, we calculated the stimulation response for any stimulation by subtracting out the mean baseline response (over a 10s period before that stimulation) from the peak response during stimulation. To test for the presence of sensory habituation, we first divided the response magnitudes for each group with respect to the response magnitude pre pairing from the same group. Subsequently, we tested for the presence of a negative linear regression between the normalized response and the number of stimulations *during pairing* (i.e. excluding pre and post pairing responses). We also tested whether this habituation effect is comparable between ChR2-eYFP and eYFP groups by testing whether there is a significant interaction effect between the effect of pairings and the effect of group on the normalized response magnitude. To test for an enhancement in the fMRI signal due to pairings in addition to a shared habituation, we tested the ratio of responses for each pairing (including pre and post pairing) between the ChR2-eYFP and eYFP groups for a positive linear regression.

